# Correlating Disordered Activation Domain Ensembles with Gene Expression Levels

**DOI:** 10.1101/2024.10.19.619222

**Authors:** Eduardo Flores, Aleah R. Camacho, Estefania Cuevas-Zepeda, Mary B. McCoy, Feng Yu, Max V. Staller, Shahar Sukenik

## Abstract

Transcription factor proteins bind to specific DNA promoter sequences and initiate gene transcription. In eukaryotes, most transcription factors contain intrinsically disordered activation domains (ADs) that regulate their transcriptional activity. Like other disordered protein regions, ADs do not have a fixed three-dimensional structure and instead exist in an ensemble of conformations. Disordered ensembles contain sequence-encoded structural preferences which are often linked to their function. We hypothesize this link exists between the structural preferences of disordered AD ensembles and their ability to induce gene expression. To test this, we used FRET microscopy to measure the ensemble dimensions of two activation domains, HIF-1α and CITED2, in live cells, and correlate this structural information with transcriptional activity. We find that point mutations that expanded the HIF-1α ensemble increased transcriptional activity, while those that compacted it reduced activity. Conversely, CITED2 showed no correlation between ensemble dimensions and activity. Our results reveal a sequence-dependent relationship between AD ensemble dimensions and their transcriptional activity.

**WHY IT MATTERS:** Transcription factors have activation domains (ADs) that bind to coactivator complexes to initiate gene transcription. Despite their key role, a comprehensive understanding of what drives their transcriptional activity has remained elusive. Efforts to understand AD activity have largely focused on their amino acid composition. In recent years, it is increasingly realized that the structural ensembles of disordered proteins contain biases that dictate their structural properties. For ADs, ensemble structures remain poorly explored, especially in relation to their activity. Here we report a mutational study of two ADs, HIF-1α and CITED2, that examines how ensemble dimensions correlate with activity. Our findings suggest that ensemble dimensions may drive activity in some ADs, and that AD ensemble dimensions can be modulated not only through mutations, but also through changes in the cellular environment.

## INTRODUCTION

Transcription factors bind to DNA and use their activation domains (ADs) to recruit the transcriptional machinery to the site of transcription, initiating the process of gene expression (1). TFs have DNA binding domains that facilitate DNA binding directly and have a well-defined structure. In contrast, ADs are intrinsically disordered (2–5). Intrinsically disordered regions (IDRs) lack a well-defined tertiary structure and instead are composed of a dynamic collection of rapidly changing conformations, collectively referred to as an ensemble (6). Recent studies have used high-throughput experiments to link amino acid composition and the ability of ADs to induce gene expression (7–9). This work has provided evidence that in many ADs, acidic residues and intrinsic disorder work together to expose hydrophobic motifs, thereby enhancing their ability to bind to coactivators. However, recent evidence suggests that the acidic exposure model may not fully account for all ADs, as some may have specific positions where single amino acid substitutions, even beyond the residues proposed by the model, can fine-tune transcriptional activation (10). Despite significant experimental progress, most research has focused primarily on the amino acid sequence of ADs, with less attention given to the role that ensemble dimensions may play in regulating activity.

While the link between AD sequence and activity has been studied extensively, few studies comprehensively explore the link between ensemble and function. We hypothesize that expanded ensemble dimensions in ADs can improve coactivator binding, resulting in higher gene expression (**Fig. 1A**). In contrast, compact AD ensembles would hinder coactivator binding, resulting in low gene expression. The acidic exposure model could be seen as a specific example within the broader concept of expansion and compaction, connecting AD sequence, structure and function.

**Fig. 1.**
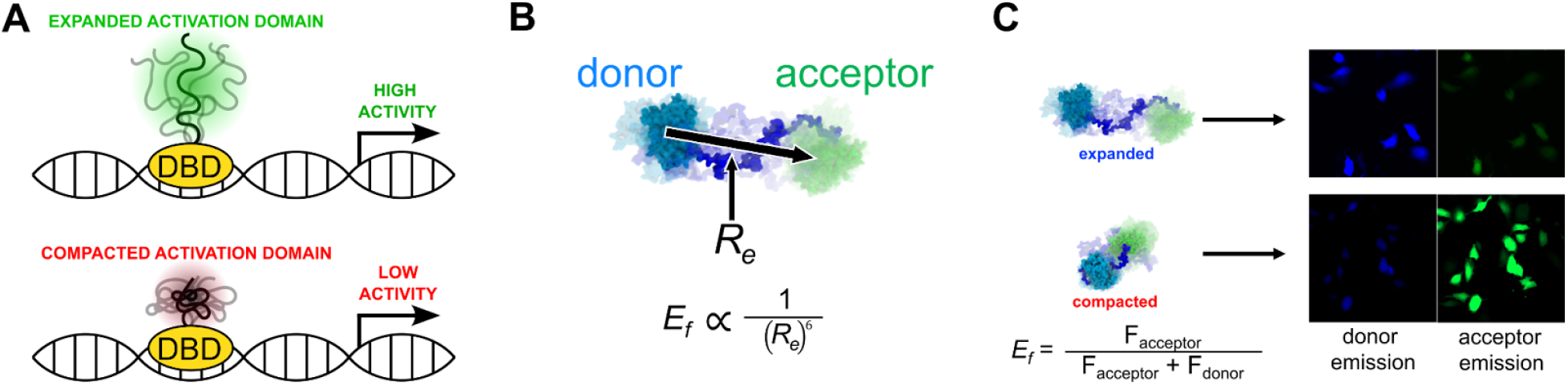
The relationship between ensemble dimensions and activity correlated using live-cell FRET. **(A)** The acidic exposure models stipulates that disordered activation domains (ADs, green and red squiggly lines) need an expanded ensemble to induce high levels of gene expression. **(B)** In the FRET constructs used here, AD sequences (in dark blue) are flanked by two fluorescent proteins - the donor, mTurqouise2 is shown in cyan and the acceptor, mNeongreen is shown in green. The distance between the fluorophore center of the fluorescent proteins, *R*_*e*_, is inversely proportional to the FRET efficiency (*E*_*f*_). **(C)** Constructs shown in (B) are transiently expressed in U-2 OS cells and their ensemble dimensions, as characterized by *E*_*f*_ in live cells using FRET microscopy. Donor and acceptor fluorescence intensities are quantified, corrected for artifacts, and used to calculate the average FRET efficiency (*E*_*f*_) over entire cells.

## MATERIALS AND METHODS

### AD activity analysis

Activation domain sequences and activity data for HIF-1α, CITED2, and their variants, as well as their standard deviations, and the p-values adjusted using a 5% false discovery rate were obtained from Staller et al. (7). Activity fold change was calculated by dividing the activity in the mutant by the activity of the wild type (WT) protein.

### Sequence and ensemble feature analysis

*R*_*e*_ prediction and sequence feature analysis was done using ALBATROSS (11) and CIDER (12) software packages respectively. Sequence features statistical analysis was done by comparing activating and deactivating sequences using an independent, two-sided t-test. *R*_*e*_ was normalized by dividing the *R*_*e*_ value of each mutant by the *R*_*e*_ value of the WT (**Fig. 1B**). All sequence features are described in (11). To determine the fraction of residues that fall under the acidic exposure model (AEM), the fraction of aromatic, leucine, and acidic residues was calculated using the `compute_residue_fractions`() function within SPARROW (https://github.com/idptools/sparrow).

### Plasmid handling

Plasmids for mammalian expression were constructed by inserting the mTurquoise2 and mNeonGreen genes into a pcDNA3.1(+) plasmid. This was done using the 5′ NdeI and 3′ XhoI restriction sites. The IDP regions were obtained from GenScript and inserted between the two fluorescent proteins (FPs) using the 5′ SacI and 3′ HindIII restriction sites. The resulting plasmids were then amplified in XL-1 Blue cells (Thermo Fisher Scientific) following the manufacturer’s protocol. The amino acid sequences of all FRET constructs, including their IDP inserts, are provided in **Table S1**.

### Mammalian cell culture and plasmid transfection

U-2 OS cells (ATCC) were cultured in T25 (Thermo Fisher) flasks using Advanced DMEM (Gibco) supplemented with 10% fetal bovine serum (Gibco) and 1% penicillin-streptomycin (Gibco). For live cell microscopy experiments, 10,000 U-2 OS cells were plated in a μ-Plate 96-well black plastic, ibidi-treated imaging plate (Ibidi) and allowed to adhere overnight (∼16 hours) before transfection. The cells were maintained at 37 °C with 5% CO_2_. Before transfection, the media was replaced with fresh, warmed DMEM. XtremeGene HP (Sigma) was used to transfect FRET construct plasmids into U-2 OS cells, following the manufacturer’s protocol. After transfection, the cells were incubated at 37 °C with 5% CO2 for 48 hours.

### Live cell microscopy

Imaging was performed using a Zeiss epifluorescent microscope equipped with a 20× objective (0.9 NA, dry). The samples were illuminated using a Colibri2 LED light engine (Zeiss), and data were collected with a duo-cam setup, which included two linked ORCA Flash v3 sCMOS cameras (Hamamatsu). The cells were imaged at an ambient temperature of 21 °C, both before and after perturbation, with exposure times of 150 ms and an LED intensity set to 7%. mTurquoise2 was excited at 430 nm (both donor and acceptor channels), and mNeonGreen was excited at 511 nm (direct acceptor channel). Emitted light was passed on to the cameras using a triple bandpass dichroic filter (467/24, 555/25, 687/145). For FRET measurements, the emitted light was split onto the two cameras using a downstream beam splitter with a 520 nm cutoff. Images were taken in PBS or HEPES buffers for mammalian cell culture.

For cell volume perturbation cells were first imaged, followed by another image after the addition of perturbation solution. To ensure thorough mixing, the solutions were pipetted up and down ten times prior to imaging. The second image was captured within approximately 25 seconds. Perturbation solutions were created by either diluting the imaging medium (1× PBS) with autoclaved deionized water to achieve hypo osmotic conditions (100 mOsm final osmotic pressure) or by adding 1 M NaCl stock solution to achieve hyperosmotic conditions (750 mOsm final osmotic pressure). Isosmotic conditions (300 mOsm) were obtained by using 1× PBS (Gibco). Before imaging, cells were rinsed once with 1× PBS and then left in 200 μl of 1× PBS (300 mOsm) prior to imaging.

### Image analysis

Images were analyzed using ImageJ(13). The images collected before and after the osmotic challenge, which contained three channels each, were stacked and aligned using the StackReg plugin with a rigid transformation(14). The aligned images were segmented based on the donor channel before perturbation, using a fixed threshold to select only pixels with intensities higher than 850 AU. The resulting mask was processed using ImageJ’s Watershed binary algorithm. Cells were selected using the Analyze Particles option in ImageJ, filtering for those with an area between 200 μm^2^ and 50,000 μm^2^ and a circularity between 0 and 0.718. The average intensity of the selected regions of interest was calculated. Cells with average direct acceptor emission intensities ranging from 1,258.9 and 7,943.2 AU were selected as lower values were at the noise levels, and higher intensities introduced artifacts and oligomerization effects(15). Cells with an absolute change in direct acceptor emission following osmotic change greater than 1,200 AU (typically cells that moved or lifted off the coverslip during measurement) were excluded.

To correct for donor bleedthrough and cross-excitation, cells were transfected with either the mTurquoise2 or mNeonGreen construct alone, as described previously(15). These cells were imaged and analyzed using the same protocol described above, and correlation plots were generated to determine the percentage of bleedthrough and cross-excitation. In the final filtering step, cells with a corrected donor/acceptor ratio that was negative or greater than 6 were removed. The analysis code is available as an ImageJ macro at the accompanying github repo for this paper (https://github.com/sukeniklab/Flores-et-al-2024).

### FRET Analysis

FRET efficiencies for perturbations and basal conditions were calculated for each segmented cell as follows:

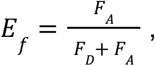

where *F*_*D*_ and *F*_*A*_ represent donor and bleed through and cross-excitation corrected acceptor emissions, respectively (**Fig. 1C**). Each emission value is the average of an entire segmented cell so that each *E*_*f*_ value represents the average FRET efficiency of a single cell. The *E*_*f*_ of all the cells in a single imaging well were used to construct a violin plot. Each well contained at least 30 cells, with some wells holding hundreds of cells. Significance in *E*_*f*_ was assessed based on the medians of all imaged wells (n) for two constructs using an independent, two-sided t-test. For osmotic perturbations, Δ*E*_*f*_ was calculated as: Δ*E*_*f*_  = *E*_*f,after*_ *-E*_*f,before*_ for all osmotic perturbations, where *E*_*f*_ is again calculated per cell, with all cells from a single well were used to create a single violin plot. Statistical analysis for these experiments also used an independent t-test to compare hypo- or hyperosmotic with isosmotic perturbation Δ*E*_*f*_ values. The number of cells measured for each construct and condition is summarized in **Table S2 and S3**.

## RESULTS

### Point mutations result in distinct ensemble profiles for HIF-1α and CITED2 activation domains

We began by comparing activity measurements published previously for HIF-1α and CITED2 that included the activity of sequences with multiple mutations (7). Point mutations applied to HIF-1α were able to both increase or decrease activity significantly compared to the wildtype (**Fig. 2A**). In contrast, CITED2 point mutations showed no significant increase in activity, and in many cases only decreased activity (**Fig. 2B**).

**Fig. 2.**
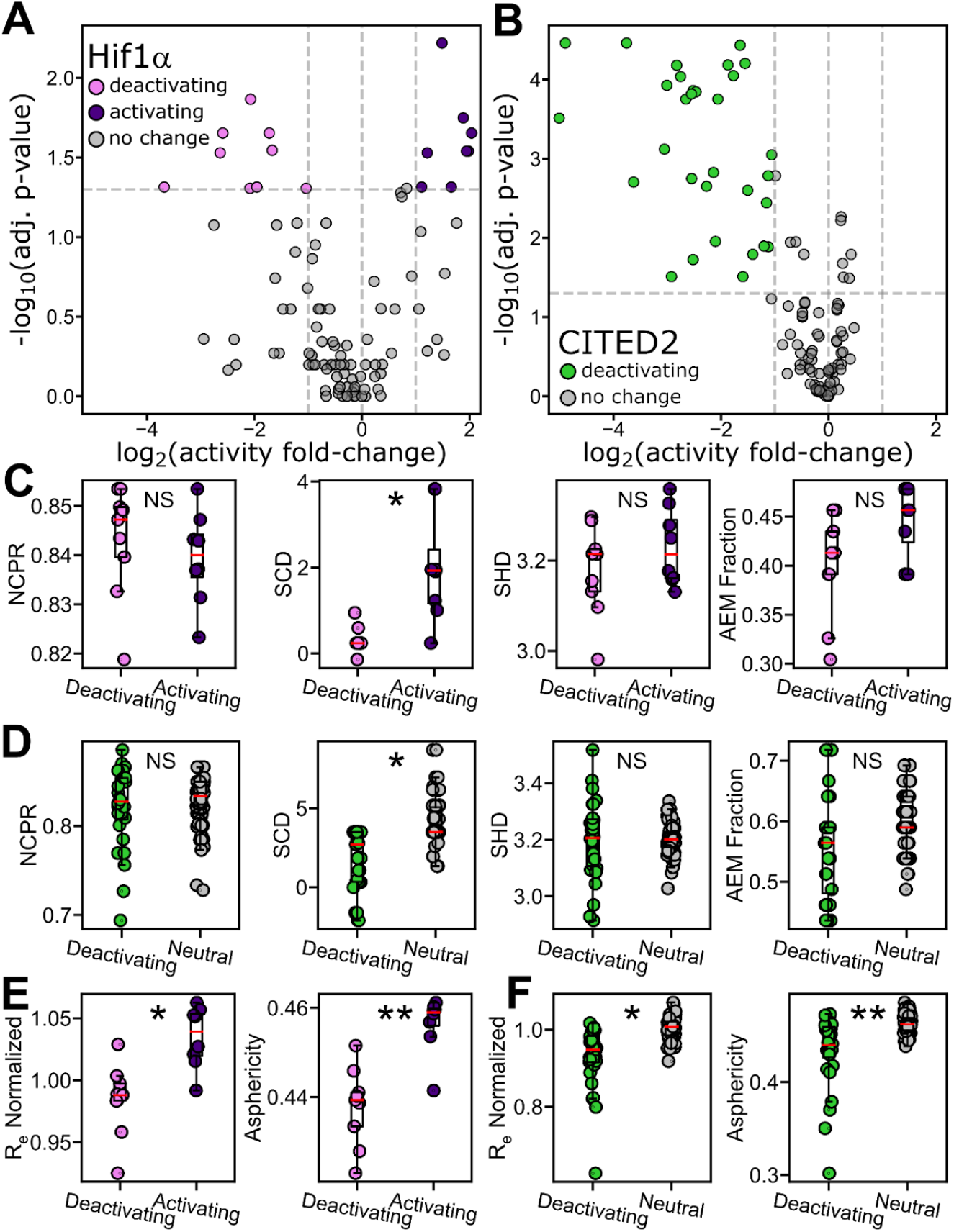
HIF-1α and CITED2 transcriptional activity levels correlated with sequence properties. **(A)** Volcano plot of HIF-1α fold change in mutant activity compared to the WT sequence. Significantly activating and deactivating mutations are shown in purple and pink symbols respectively (where p-value adjusted with a 5% FDR is lower than 0.05). **(B)** Volcano plot of CITED2 showing mutants that significantly decrease activity (green). **(C)** HIF-1α sequence features of significantly activating and deactivating HIF-1α mutants. (D) CITED2 sequence features significantly activating mutants and those with no effects. Statistical significance in (C) and (D) is denoted through asterisks (*, **, and *** denote p-values of < 0.01, 0.001, and 0.0001, respectively, using an independent, two-sided t-test). **(E, F)** Normalized *R*_*e*_ ALBATROSS and asphericity predictions of HIF-1α and CITED2 mutants respectively.

We next wanted to see if we could find sequence properties that would correlate with these changes in activity. To do this we calculate the sequence charge decoration (SCD) (16), sequence hydropathy decoration (SHD) (17), and the fraction of residues participating in the acidic exposure model (Trp/Asp/Glu, AEM fraction) (**Fig. 2C, D**). These were selected to explore whether charge-related (NCPR, SCD) or non-charge related (SHD) properties could explain the differences between activating and deactivating sequences. Lastly, we measured the fraction of amino acids that fall within the AEM to determine if this metric revealed any significant differences between the sequences (**Fig. S1**). In HIF-1α, activating sequences had a significantly higher SCD when compared to deactivating sequences (**Figs. 2C**). All other parameters we tested showed no significant difference (**Figs. 2C, S1**). CITED2 had no activating mutations, so instead we used mutations that had no significant effect on activity compared to WT (|log_2_ (FC)| < 1 or -log_10_ adjusted p-value < 1.3). Here again, the only sequence feature that showed a significant difference between deactivating and neutral sequences was the SCD (**Fig. 2D, S2**). This result led us to explore whether differences might be more significant in the structural features of the ensemble.

We predict ensemble features using ALBATROSS (11). Unlike sequence properties, predicted ensemble features showed significant correlations with activity. HIF-1α showed a statistically significant difference between activating and deactivating sequences in the normalized predicted end-to-end distance (*R*_*e*_) and asphericity (**Fig. 2E**). Similarly, CITED2 displayed significant differences between deactivating and neutral sequences for these same ensemble features (**Fig. 2F**). Specifically, mutations predicted to expand ensemble dimensions were associated with increased activity, while those predicted to cause compaction were linked to decreased activity. In contrast, CITED2 showed that most mutations resulted in a decrease in activity while having no significant effect on *R*_*e*_ (**Figs. 2F, S2**). A growing body of work links the ensemble dimensions of disordered regions to their function (6, 18, 19). This, together with the correlations between ensemble structure and activity for ADs we found, motivated us to experimentally explore the link between AD ensemble structure and function directly.

### Live cell FRET reveals that HIF-1α, but not CITED2, displays structural plasticity

To establish the link between HIF-1α and CITED2 AD ensembles and their activity, we selected a subset of mutations whose activities were measured (**Fig. 3A**). We reasoned that variants where hydrophilic and hydrophobic residues were mutated to aromatics would bring distal regions of the protein closer together through *π*-stacking interactions, compacting the ensemble (20, 21). We also reasoned that replacing all positively charged residues with negatively charged residues would introduce electrostatic repulsive forces with existing negative charges that would expand the sequence (22). Lastly, we wanted to see what structural role positive charges, hydrophobics and aromatics played when absent from the WT CITED2 (FL>A_NT_ and RK>A_CT_).

**Fig. 3.**
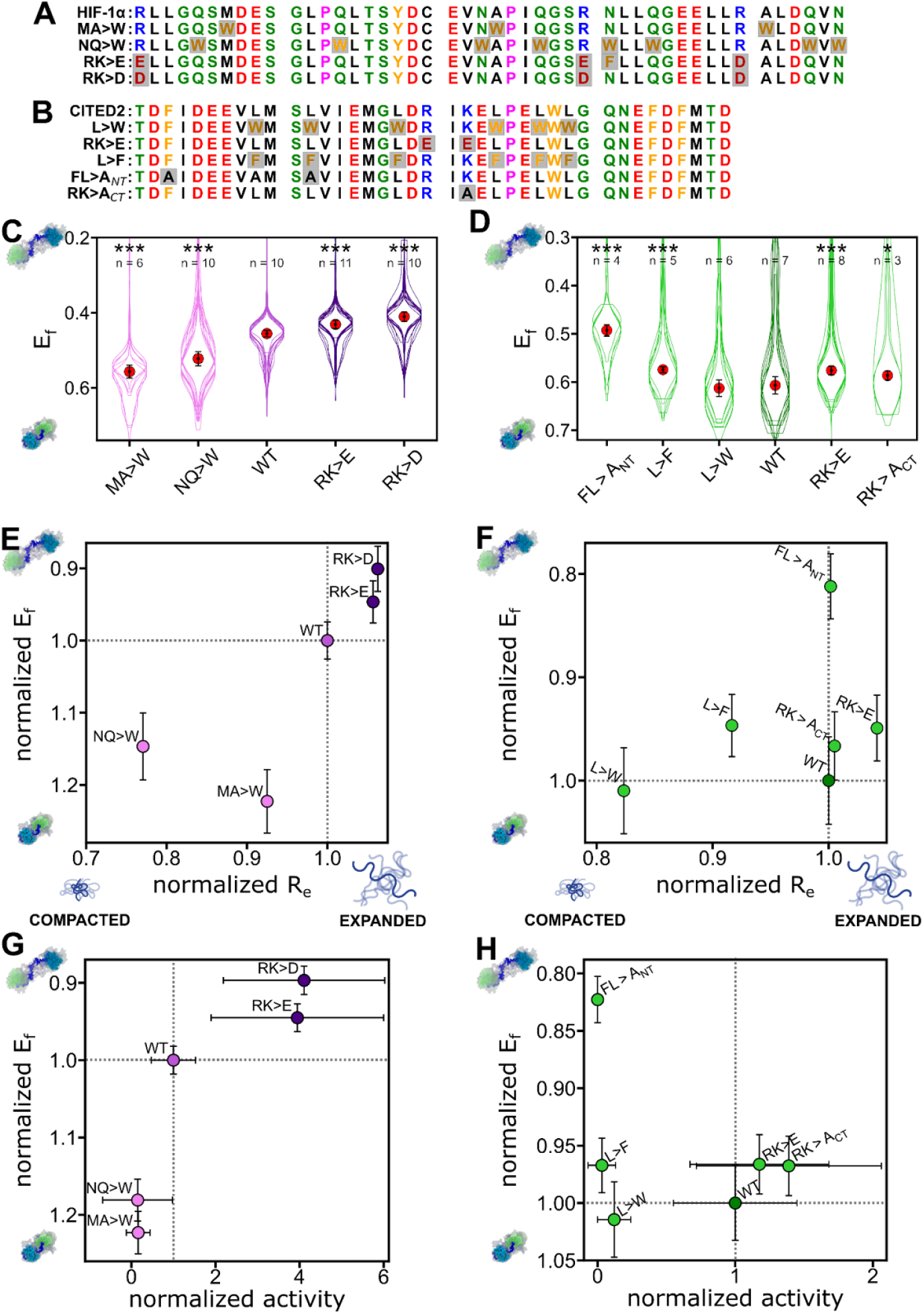
Ensemble dimensions of HIF-1α measured in live cells. **(A)** WT and mutant HIF-1α sequences. Mutated residues are highlighted in light gray. **(B)** Same as (A) for CITED2. **(C)** FRET efficiencies (*E*_*f*_) in live cells for HIF-1α and its mutants. Each overlaid violin plot is obtained from an individual experiment done in a single imaging well and contains at least 30 cells. The red point is the average of the medians of n wells, where n is shown at the top, and the black whiskers represent the standard deviation of the medians. Asterisks denote significant difference from the WT (*, **, and *** denote p-values of < 0.01, 0.001, and 0.0001, respectively). **(D)** same as (C) for CITED2. **(E)** Normalized *vs* predicted end-to-end distance normalized by the WT end-to-end distance. (**F**) same as (E) for CITED2. **(G)** Normalized *E*_*f*_ of HIF-1α mutants normalized *E*_*f*_ *vs* activity normalized to that of the WT. (**G**) same as (F) for CITED2.

To measure the ensemble dimensions of the selected mutants experimentally we used live-cell FRET microscopy (15). Ensemble dimensions are measured through the FRET efficiency, E_f_, between a donor and an acceptor fluorescent protein (**Fig. 1B**), where a high efficiency denotes a compact ensemble and a low efficiency denotes an expanded ensemble (**Fig. 1C**). Our measurements reveal that the selected HIF-1α mutants behaved as predicted: Sequences with aromatic mutations resulted in more compact ensembles than the WT sequence as indicated by a higher *E*_*f*_. HIF-1α sequences that introduced negatively charged residues expanded the ensemble as shown by a decrease in *E*_*f*_ compared to the wildtype (**Fig. 3C**). We also correlated the resulting *E*_*f*_ of mutations with the predicted *R*_*e*_ from ALBATROSS. All HIF-1α sequences showed a correlation (pearson’s r = 0.78) between the average median *E*_*f*_ and the predicted Re, suggesting that our simulations align well with live-cell FRET measurements, even with the attached flanking fluorescent proteins (**Fig 3E**).

In contrast, CITED2 variants displayed some unexpected behavior with respect to ensemble dimensions. First, variants with insertion of aromatics (L>W) did not compact the ensemble as we hypothesized. Instead, these variants maintained similar ensemble dimensions to the WT (L>W) or, even more surprisingly, led to increased expansion (L>F) (**Fig. 3D**). Replacing positively charged lysine and arginine with negatively charged glutamate (RK>E) led to an expansion of the ensemble, likely attributed to the introduction of repulsive electrostatic forces. Substituting aromatic phenylalanine and leucine residues with alanine (FL>A_NT_) led to the ensemble’s expansion, consistent with the removal of hydrophobic and aromatic residues (**Fig. 3D**) Lastly, substitution of positively charged residues with alanine in the C-terminus (RK>A_CT_) resulted in a small expansion in its ensemble dimensions. Unlike HIF-1α, *E*_*f*_ for CITED2 had poor correlation with *R*_*e*_ (Pearson’s r = -0.08), which was predicted to be more variable, indicating the possibility of IDR:FP interactions (**Fig. 3F**).

To finally check if ensemble dimensions correlated with activity, we plotted the average median *E*_*f*_ against the activity measured for the same AD. Both the average median *E*_*f*_ and activity were normalized to the associated WT construct. We observed a positive correlation (pearson’s r = 0.90) between the normalized average median *E*_*f*_ and the normalized activity in HIF-1α sequences, indicating that an expanded ensemble facilitates higher HIF-1α activation, while a compact HIF-1α ensemble would hinder activity (**Fig. 3G**). CITED2, on the other hand, showed a contrasting profile. Mutations either completely abolished transcriptional activity, even when ensemble dimensions remained unchanged or expanded, or maintained activity at levels similar to the WT, showing little correlation with ensemble dimensions (Pearson’s r = 0.47) (**Fig. 3H**).

The HIF-1α mutants support the acidic exposure model and link ensemble dimensions for AD function. The HIF-1α mutants that exhibit expanded ensemble dimensions often feature acidic residues, and the repulsion between those could drive ensemble expansion (RK>E, RK>D). Conversely, HIF-1α mutants that are ‘collapsed’ feature mutations into aromatic residues (NQ>W, MA>W) which are known to form attractive interactions and can thus collapse the ensemble.

In contrast, we hypothesize that the WT CITED2 ensemble is structurally rigid, which makes further expansion through sequence modifications difficult. Structurally, only the FL>A_NT_ mutant resulted in a compact ensemble, while the other mutants maintained similar ensemble dimensions when compared to the WT. It’s important to note that the FL>A_NT_ mutant occurs in a region previously characterized as activity-driving(7). Thus, decrease in activity may not be related to changes in ensemble dimensions but rather a result of changes in binding affinity or other mechanistic functions.

### Compact high-activity sequences are structurally sensitive to osmotic perturbations

Our results show that ensemble dimensions can be modulated through changes in sequence, but past work indicates that they can also be modulated from changes in the cellular environment (15, 23). If the link between ensemble dimensions and activation domain activity/function holds true, it might mean that the cellular environment could modulate AD ensembles, which acts as a previously unappreciated mechanism for regulating gene expression. To see if this hypothesis is a possibility, we decided to test the response of ADs to cellular volume change perturbations. Although these perturbations may not be directly relevant to the physiological activities of CITED2 or HIF-1α, they still provide insights into the structural plasticity of the sequence in a changing cellular environment.

Osmotic challenges act to rapidly and reproducibly modulate the volume of the cell (24). Cellular volume changes occur routinely during cell cycle and change the composition of all solute molecules inside the cell (25, 26). Our previous work has shown that some disordered regions could be sensitive to osmotic changes (15). Thus, their effect on ensemble structure could also be meaningful to AD activity. To test the structural sensitivity of AD ensembles to changes in the cellular environment, we measured changes in FRET efficiency 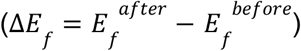 of the HIF-1α and CITED2 ADs inside the cell as it is subjected to hypo- or hyper-osmotic challenge. We also measured the response of a control sequence composed of a sequence composed of 24 Gly-Ser repeats (GS24) which acts as a homopolymeric point of reference (15). This point of reference allows us to gauge if the sequence displays sensitivity different than that of a homopolymer to changes in cell volume (15, 23).

HIF-1α and CITED2 show opposite responses to our osmotic challenges. Hyposmotic pressure did not significantly impact the ensemble dimensions of HIF-1α when compared to the isosmotic control (**Fig. 4A**). In contrast, a hyperosmotic challenge significantly increased FRET efficiency for HIF-1α (Δ*E*_*f*_ > 0) compared to the isosmotic control, indicating a compaction of the sequence. CITED2 compacted under hyposmotic stress as indicated by an increase in FRET efficiency (Δ*E*_*f*_ > 0) (**Fig. 4B**). When exposed to the same hyperosmotic challenge, CITED2 exhibited ensemble expansion (Δ*E*_*f*_ < 0). Here we use a GS24 (48 total residues) glycine-serine repeat homopolymer for comparison (**Fig. 4C**). We observe that the homopolymer responds similarly to HIF-1α. Overall, CITED2 is more sensitive to the cellular environment than either HIF-1α or the GS24 linker, and the effect of osmotic changes on the CITED2 ensemble is significantly larger than that observed from most of its mutations studied here.

**Fig. 4.**
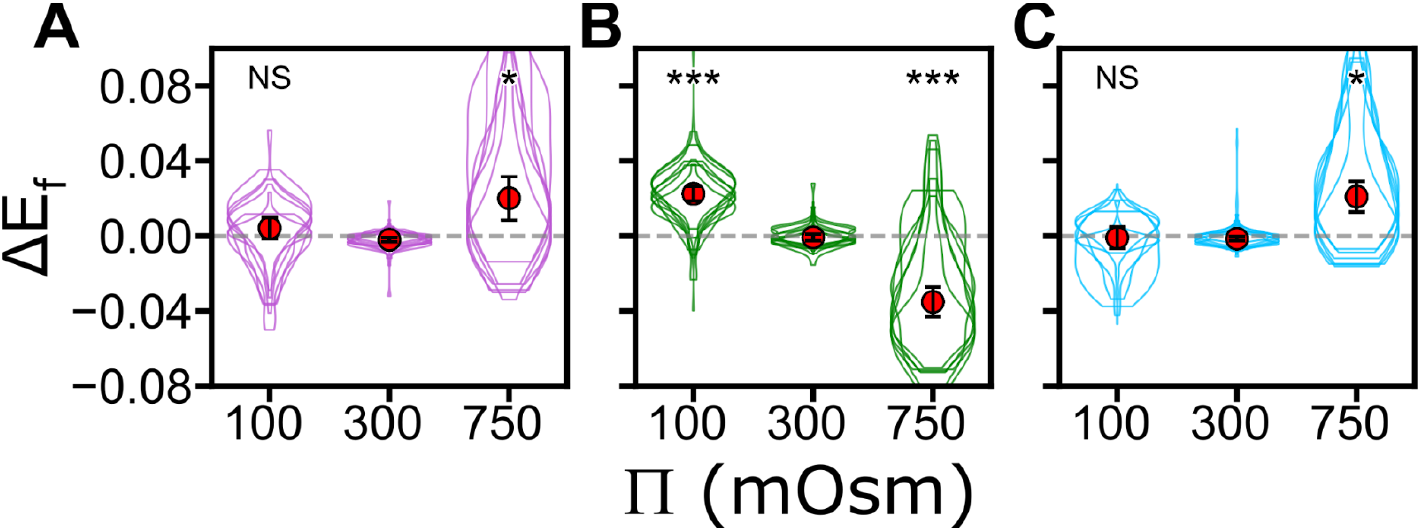
Live-cell characterization of ensemble sensitivity in response to osmotic stress. Change in FRET efficiency following osmotic preturbations, *ΔE*_*f*,_ in response to osmotic stress for HIF-1α (A), CITED2 (**B**), and GS24 (**C**). Violin plots show hyposmotic (100 mOsm), isosmotic (300 mOsm) and hyperosmotic (750 mOsm) stress. Overlaid violin plots represent multiple well repeats. Red dot denotes the average *ΔE*_*f*_ median value for the sequence along with its propagated error. Significant differences in *ΔE*_*f*_ when compared to different osmotic stress is denoted with an asterisk (*, **, and *** denote p-values < 0.01, 0.001, and 0.0001, respectively using a double-sided independent t-test).

Our findings substantiate the elusive characterization of ADs: what is true for some, might not be true for others. Given that ADs are largely disordered, we aimed to shed light on the role that ensemble structure plays in their activity. For HIF-1α, our study shows that ensemble dimensions correlate with activity. Specifically, expanded ensembles showed a higher HIF-1α gene expression activity, while compact ensembles showed lower activity levels. It has previously been suggested (27–29), though recently contended (30), that an expanded protein region enhances DNA binding by facilitating access through transient interactions. For ADs, this expanded ensemble could result in a high capture radius for the IDR, enhancing its ability to bind transcriptional coactivators necessary for transcription initiation.

For CITED2, the emerging picture is more complicated because mutations did little to change ensemble dimensions, and no mutations were found that increase the activity of the construct. We therefore propose that ADs like CITED2, in which the ensemble is less prone to changes through mutations, can be rather stable in the face of mutations. We emphasize that our findings are specific to the two activation domains studied and may not be applicable to all activation domains. Still, a common characteristic among ADs is that they are intrinsically disordered which makes them exposed to the cellular environment.

Our results contain some limitations worth considering. One key observation is the weak correlations between CITED2’s ensemble dimension and predicted *R*_*e*_ in contrast to HIF-1α. To investigate this discrepancy, we analyzed the sequence and ensemble features of their respective variants. Of all the metrics, only the fraction of charged residues was significantly larger in CITED2 compared to HIF-1α. A higher concentration of charged residues could lead to stronger interactions with the flanking fluorophores, which on their own carry some charges(15). This could potentially affect the observed *E*_*f*_. In this case, further studies of CITED2 mutations in the absence of the fluorescent protein labels might show a more flexible ensemble. Nonetheless, activity measurements, which were done without the FPs, show that no mutant had an increase in activity.

One additional finding in this work is that perturbations to cell volume had a much greater effect on CITED2 ensemble dimensions in comparison to HIF-1α or its GS24 control. Previous work has shown how changes in cell volume not only affect cell fate, but also TF function (31, 32). Further research is needed to determine whether the changes in ensemble dimensions caused by osmotic stress also correlate directly with activity. Based on our findings, we propose that the cellular environment may be a novel mechanism for regulating activity through changes in AD ensemble dimensions. It is interesting to note that the AD which displayed activity tuned by ensemble dimensions showed little sensitivity, while those whose activity was not easily tunable by mutations showed a larger sensitivity. This may indicate mutually exclusive mechanisms by which AD function is carried out.

## ACKNOWLEDGEMENTS

Research reported in this publication was supported by the NIH under award R35GM150813 to M.V.S. and R35GM137926 to S.S.

## AUTHOR CONTRIBUTIONS

S.S., E.F., M.V.S., and F.Y. conceptualized the project. E.F. designed and performed all live cell experiments, analysis, and figures, assisted by A.R.C, E. C. -Z., and M.B.M. All activity data was provided by M.V.S. S.S., M.V.S., and E.F. wrote the paper.

## COMPETING INTERESTS

The authors declare no competing interests.

## SUPPLEMENTAL INFORMATION

*Supplemental figures* S1 and S2 show analyses of all sequence and ensemble features for all constructs in Fig. 2A, B, respectively. *Table S1 -* Shows the amino acid sequence, activity, and standard deviation associated for each HIF-1α and CITED2 mutant. The amino acid sequence with the FRET pair sequence is also included. *Table S2 -* Displays the total number of cells present in each well (within a 96-well plate) for each mutant. This is performed separately for both HIF-1α and CITED2 mutants. *Table S3 -* Shows the total cell count in each well of a 96-well plate for each osmotic perturbation and mutant. This is performed separately for both HIF-1α and CITED2 mutants. All data used to produce the figures as well as scripts are available at https://github.com/sukeniklab/Flores-et-al-2024.

## Notes

https://github.com/sukeniklab/Flores-et-al-2024

### Competing Interest Statement

The authors have declared no competing interest.

