## Supplementary material for "Correlating Disordered Activation Domain Ensembles with Gene Expression Levels": Figures S1-2

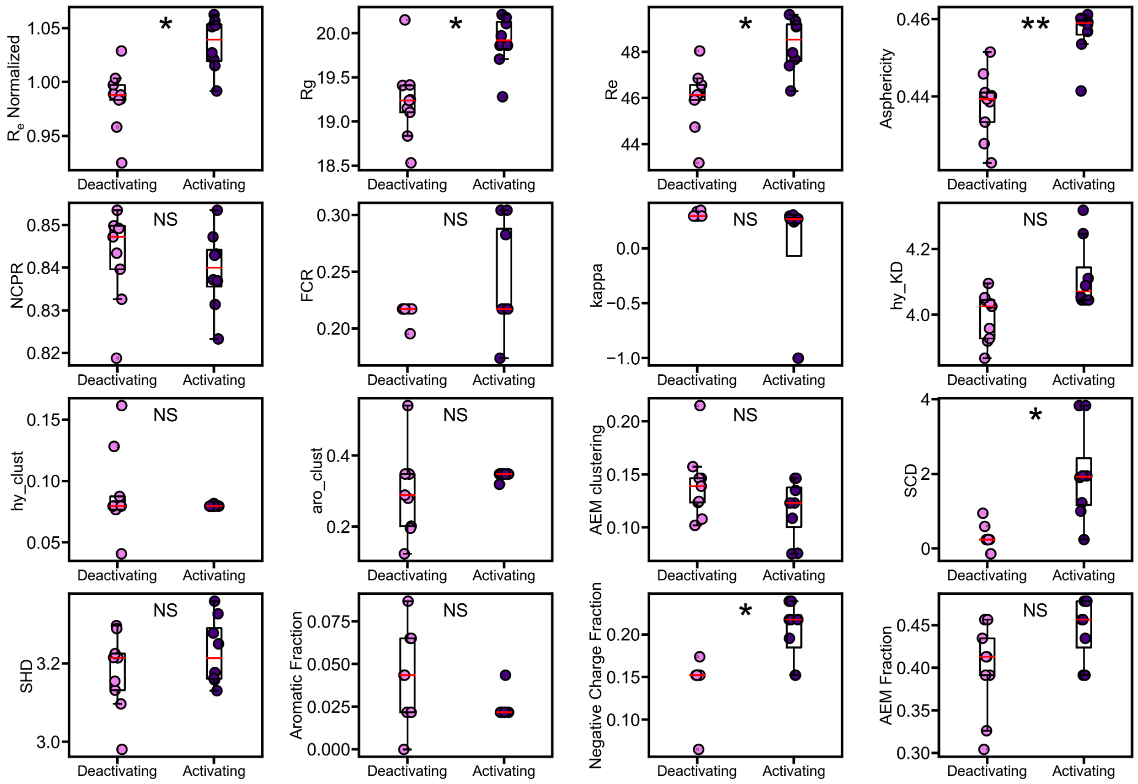


**FIGURE S1**. **Sequence features of HIF-1α.** For HIF-1α, only the significantly activating and deactivating mutants are considered in the analysis. Each feature on the y axis indicates the following: **Asphericity**: A shape descriptor that quantifies how elongated or spherical the ensemble is, with higher values indicating greater asymmetry. **hy_KD (Kyte-Doolittle Hydropathy Index)**, **hy_clust (Hydrophobic Clustering)**, **aro_clust (Aromatic Clustering)**, **AEM clustering (Acidic and Electrostatic Model Clustering)**: Identifies regions of acidic, aromatic, and leucine residues as described by the Acidic Exposure model. This is followed by how clustered these residues are. **SCD (Solvent-Accessible Charged Density)**: A measure of how exposed charged residues are to the solvent, influencing solubility and interactions with other molecules. **SHD (Solvent-Accessible Hydrophobic Density)**: A measure of how much hydrophobic surface area is exposed to the solvent, impacting the protein's stability and interactions. **AEM Fraction (Acidic and Electrostatic Model Fraction)**: The fraction of acidic, aromatic, and leucine residues as described by the Acidic Exposure model. Statistical significance between the different categories (activating, deactivating, and neutral) for each sequence feature is indicated using asterisks: * for p-values < 0.01, ** for p-values < 0.001, and *** for p-values < 0.0001. These p-values are calculated using an independent, two-sided t-test. Additionally, the normalized $R_e$ values from the ALBATROSS predictions for HIF-1α and asphericity predictions for CITED2 mutants are also included in the analysis.

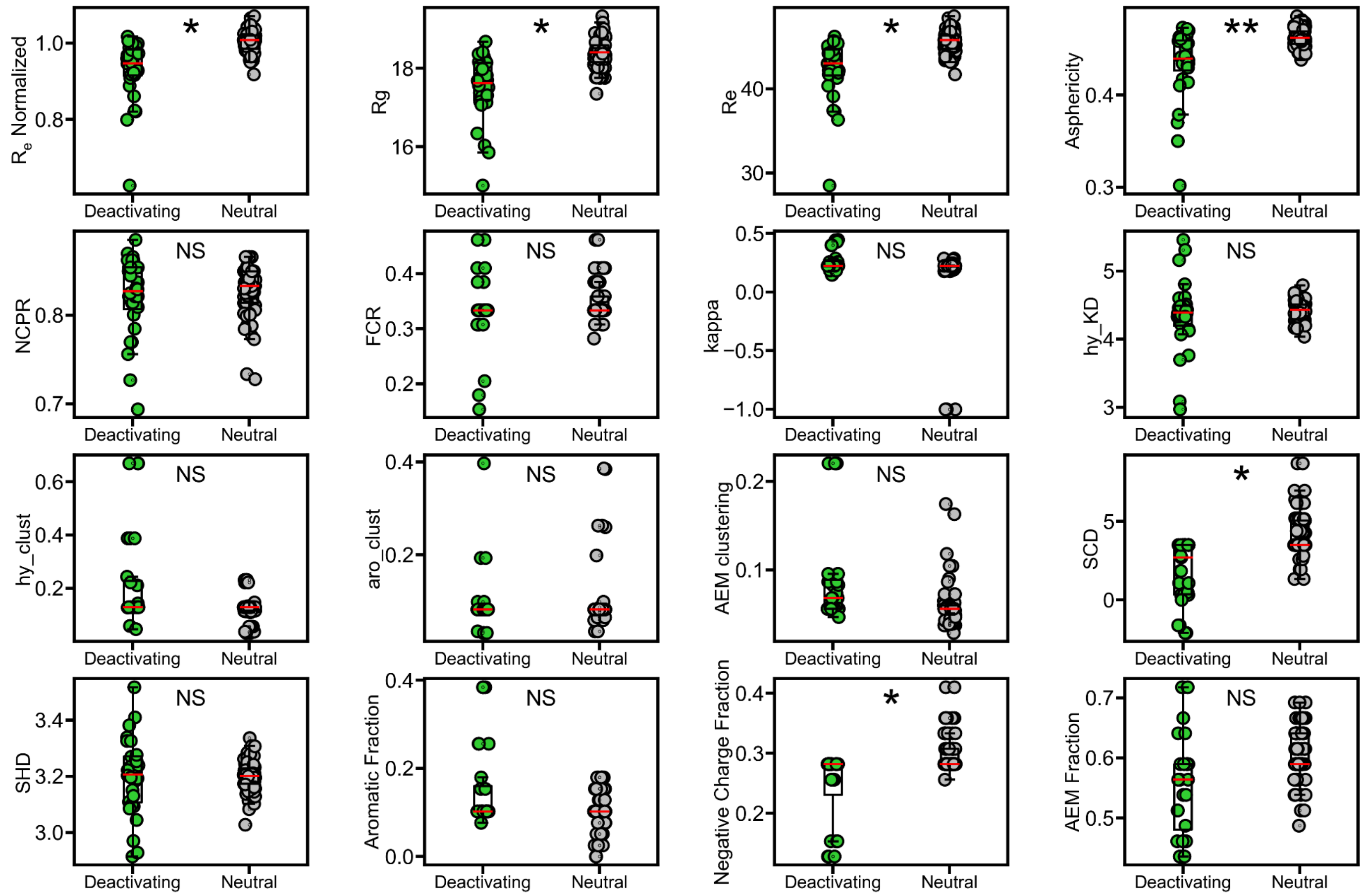


**FIGURE S2**. **Sequence features of CITED2 mutants.** In contrast, for CITED2, sequence features are compared between deactivating and neutral (gray) sequences. Y-axes are the same as in **Fig. S1**.
